# Spontaneous Neural Encoding of Social Network Position

**DOI:** 10.1101/098988

**Authors:** Carolyn M. Parkinson, Adam M. Kleinbaum, Thalia Wheatley

**Affiliations:** Department of Psychology, University of California, Los Angeles, 1285 Franz Hall, Box 951563, Los Angeles, California, USA 90095; Tuck School of Business, Dartmouth College, 100 Tuck Hall, Hanover, New Hampshire, USA 03755; Department of Psychological and Brain Sciences, Dartmouth College, 6207 Moore Hall, Hanover, New Hampshire, USA 03755

**Keywords:** social network analysis, social perception, functional MRI, multi-voxel pattern analysis

## Abstract

Humans form complex social networks that include numerous non-reproductive bonds with non-kin. Navigating these networks presents a considerable cognitive challenge thought to have comprised a driving force in human brain evolution. Yet, little is known about how and to what extent the human brain encodes the structure of the social networks in which it is embedded. By combining social network analysis and multi-voxel pattern analysis of functional magnetic resonance imaging (fMRI) data, we show that social network information about direct relationships, bonds between third parties, and aspects of the broader network topology is accurately perceived and automatically activated upon seeing a familiar other.

---

Unlike many other species that enact social behavior in loose aggregations (e.g., swarms, herds), humans form groups comprised of many long-term, intense, non-reproductive bonds with non-kin^1^. The cognitive demands of navigating such groups are thought to have comprised a driving force in human brain evolution^2^. Yet, little is known about how and to what extent the human brain encodes the structure of the social networks in which it is embedded. Here, we characterized the social network of an academic cohort (*N*=277), a subset of whom (*N*=21) completed a functional magnetic resonance imaging (fMRI) study involving viewing videos of individuals who varied in terms of “degrees of separation” from themselves (social distance), the extent to which they are well-connected to well-connected others (eigenvector centrality–EC), and the extent to which they connect otherwise unconnected individuals (brokerage). Understanding these aspects of others’ social network positions requires tracking not only direct relationships, but also bonds between third parties and the broader network topology. Pairing network data with multi-voxel pattern analysis, we show that social network position information is both accurately perceived and spontaneously activated upon encountering familiar individuals. These findings elucidate how the human brain encodes the structure of its social world, and underscore the importance of integrating an understanding of social networks into the study of social perception.

Relationships are intrinsic to human behavior. Everyday interactions are shaped not only by our own relationships, but also by knowledge of bonds between third parties and the broader social networks in which we are embedded. Well-connected individuals can effectively threaten or bolster one’s reputation^3^, those who bridge otherwise disparate groups can efficiently seek and spread information^4^, and knowledge of mutual ties influences information-sharing and trust^5^. Human social intelligence rests, in part, on a calculus that inheres in an understanding of social network structure.

Is knowledge about others’ social network positions activated only when explicit goals require it, or spontaneously, whenever we encounter familiar individuals? It may be efficient to process such information only when our goals require it (e.g., determining how to obtain information; forecasting the repercussions of a social misstep). Alternatively, it may be beneficial to activate such knowledge spontaneously when encountering others, given the importance of social network position to many aspects of behavior and to impressions of status and competence^3^,^6^. Humans spontaneously register a great deal of information when perceiving other people (e.g., intentions, traits, emotions^7^,^8^), presumably to facilitate appropriate, beneficial social interactions. Thus, the brain may run several social “daemons”– efficient, background processes that spontaneously register information useful for predicting the social repercussions of potential actions, and, more broadly, to inform cognition and behavior.

To test whether the brain spontaneously encodes the social network positions of familiar others, we scanned (fMRI) members of a real world social network (see Fig. 1; Methods) as they viewed brief videos of 12 classmates (Fig. 2). The only task was to indicate when the same video was presented twice in a row (see Methods) in order to ensure attention without any instructions to retrieve social relationship knowledge, or person knowledge more generally. Therefore, we consider any social network position information encoded while participants perform this task to be retrieved spontaneously (i.e., without instruction).

**Figure 1.**
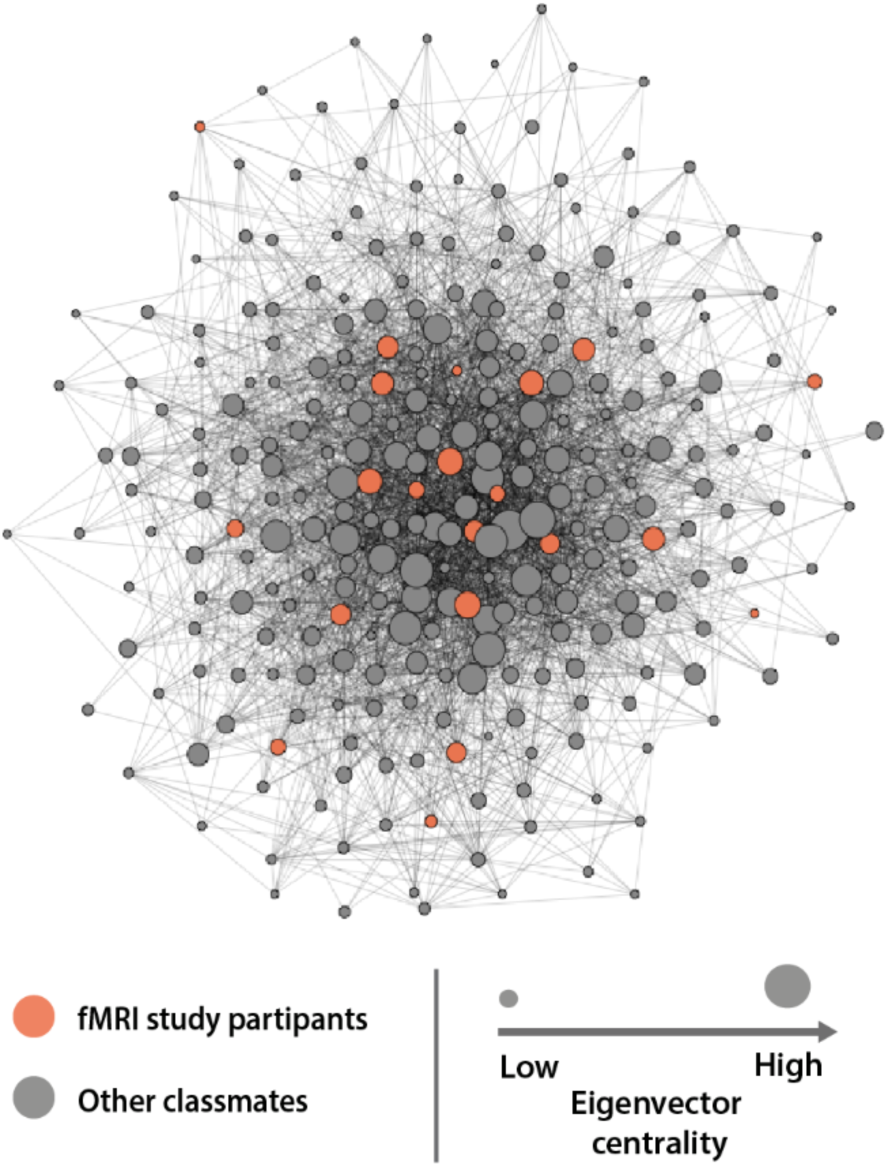
Social network characterization. The social network of a first-year cohort of MBA students was reconstructed based on responses to online questionnaires administered to all members of the class (*N* = 275; 99.3% response rate). Nodes indicate students; lines indicate reported social ties between them. For ease of visualization, only mutually reported social ties are illustrated. A subset of these students participated in an fMRI study. Orange nodes indicate fMRI study participants; gray nodes denote other members of the graduate program. Node size is proportional to eigenvector centrality.

**Figure 2.**
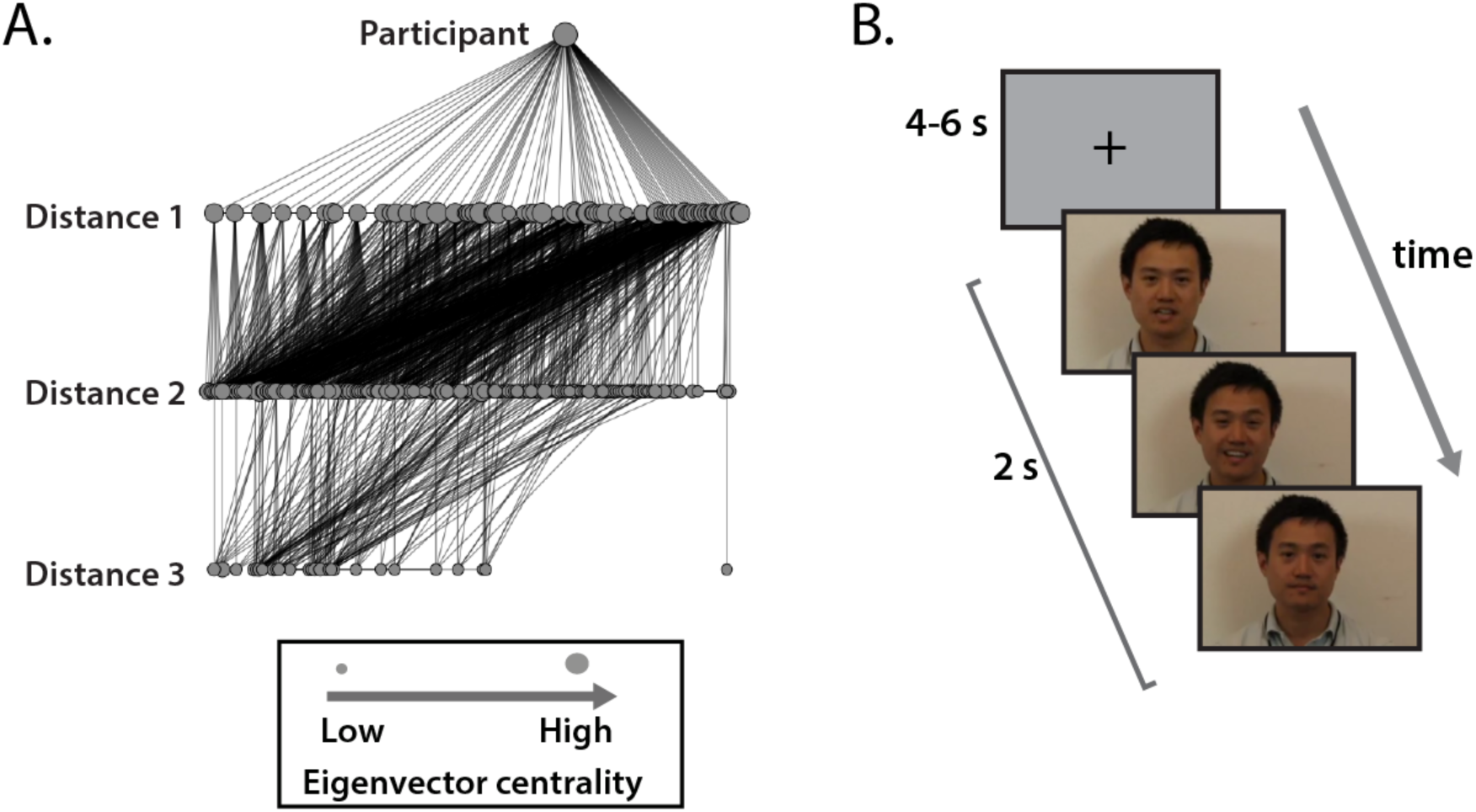
Stimulus set construction and paradigm for neuroimaging study. **(A)** The geodesic distance between each fMRI study participant and every other student in the network was characterized. An alternative visualization of the network is shown in which nodes are organized into horizontal layers according to distance from a particular participant. Each participant’s stimulus set was comprised of 12 of his or her classmates: the two lowest and two highest eigenvector centrality individuals at distances of one, two, and three from the participant in the network (e.g., the classmates signified by the two smallest and two largest nodes within each layer in **(A)**). **(B)** During the fMRI study, participants viewed brief (2 s) videos of the 12 individuals in their stimulus sets separated by 4-6 s of fixation. In order to maintain attention, a one-back task was used (i.e., participants were instructed to use a button press to indicate when an identical video was presented twice in a row). Frames from this participant’s video clip are reproduced with permission from the individual.

Each classmate in each participant’s stimulus set was characterized according to three metrics derived from the social network data: geodesic social distance from the participant; EC; and constraint, an inverse measure of brokerage. Geodesic social distance refers to the minimum number of intermediary social ties required to connect two individuals. EC is a prestige-based centrality metric that considers not only how many connections a given individual has, but also the centralities characterizing each contact^9^. High EC implies that an individual is well-connected to well-connected others; low EC implies that an individual has few friends who tend to be unpopular. Prestige-based centrality metrics are particularly useful for characterizing social status, given that being named as a friend by a popular individual should increase one’s sociometric status (i.e., the extent to which someone is liked by others) more than being named by someone less popular^9^. Individuals who connect others who would not otherwise be connected occupy network positions low in constraint, and have the capacity to serve as “brokers” of resources (e.g., information) in the network. Because of the structure of their local social ties, brokers can coordinate behavior and translate information across structural holes in networks^4^.

To probe for the spontaneous encoding of social network position information, we used representational similarity analysis (RSA), which distills fMRI response patterns into representational dissimilarity matrices (RDMs) that indicate the degree to which particular brain regions distinguish between sets of stimuli or mental states^10^. Because RDMs are abstracted away from the spatial layout of neuroimaging data (i.e., they are indexed by experimental condition; Fig. 3), RSA affords the evaluation of the degree to which similarity structures contained in particular brain regions reflect those of data acquired using other modalities of measurement or computational models^10^ (here, the social network data). Specifically, in the current study, we used a general linear model (GLM) decomposition searchlight approach^11^. Neural RDMs were iteratively extracted within 9-mm radius spheres centered at each point in each participant’s brain. Within each participant, each local neural RDM was modeled as a weighted combination of RDMs based on properties of the social network positions of the individuals in that participant’s stimulus set (Fig. 3). Using this technique, participants’ brains were mapped in terms of the degree to which the representational content of local neural responses to familiar others could be explained by those individuals’ positions in their social network, and in terms of where information about specific social network position characteristic was carried reliably across participants (Fig. 4).

**Figure 3.**
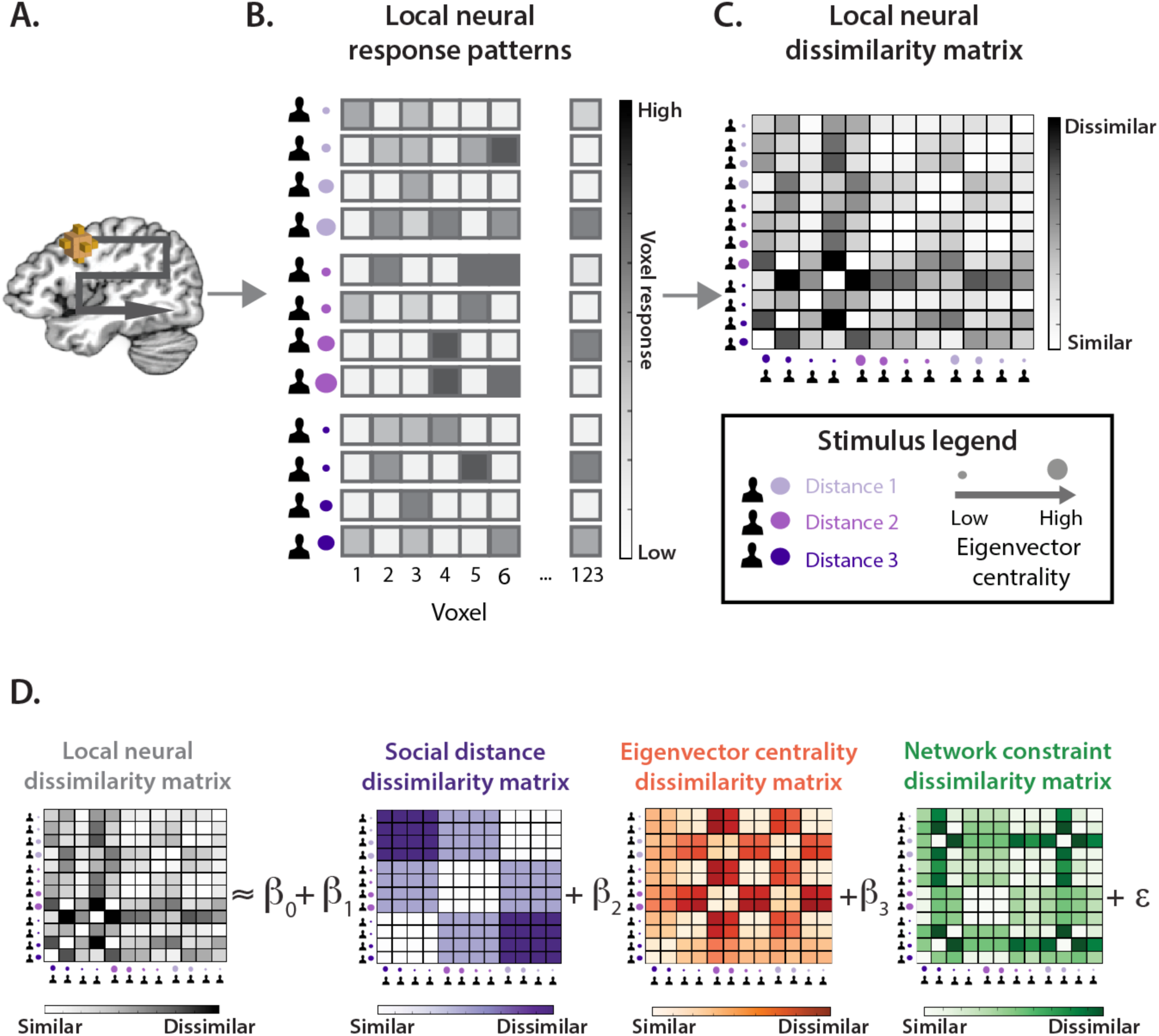
GLM decomposition searchlight. **(A)** A spherical searchlight was moved throughout each participant’s brain. **(B)** At each point in the brain, distributed patterns of neural responses to each individual in the participant’s stimulus were extracted within a 9mm radius sphere centered on that point. **(C)** At each searchlight center, a neural RDM was generated based on pairwise correlation distances between local neural response patterns to each classmate in the participant’s stimulus set. **(D)** Each local neural RDM was modeled as a weighted combination of RDMs constructed based on the pairwise Euclidean distances (i.e., the absolute value of numerical differences) between individuals in each participant’s stimulus set in terms of social distance, eigenvector centrality, and network constraint.

**Figure 4.**
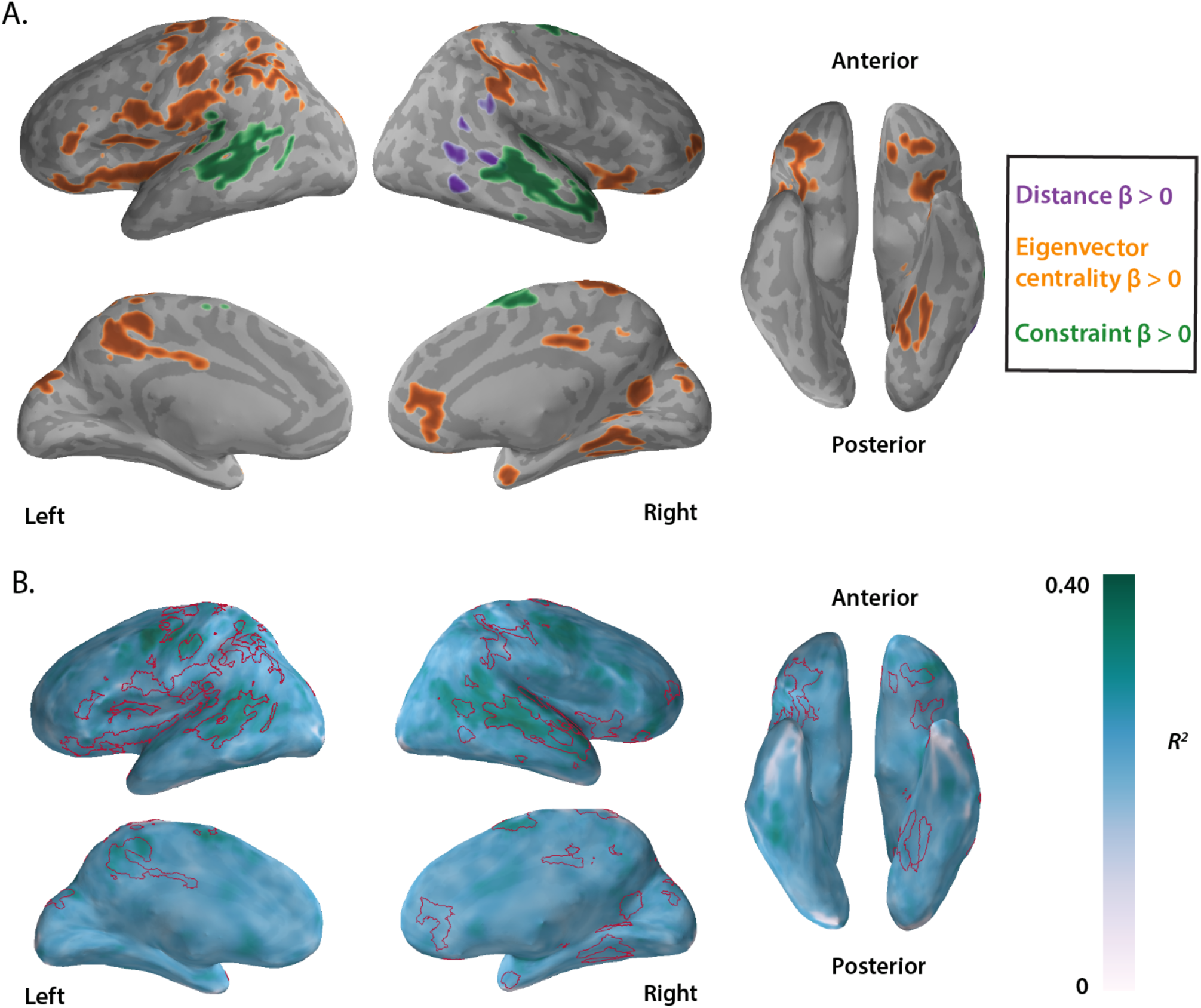
Neural encoding of social network position. **(A)** Distinct brain regions encode different properties of peers’ social network positions (social distance = purple; eigenvector centrality = orange; constraint = green). Beta values indicate the extent to which the information contained in local multi-voxel response patterns to participants’ classmates could be predicted based on properties of those individuals’ social network positions; *p* < .05, FWE-corrected. **(B)** The *R^2^* value corresponding to the GLM decomposition performed at each searchlight center indicates the extent to which the information contained in local multi-voxel response patterns can be explained by the social network positions of the classmates being viewed. Voxel-wise *R^2^* values averaged across subjects are depicted; red contours indicate clusters of voxels that reliably signaled one or more of the tested aspects of social network position across participants. Results are projected onto a cortical surface model of the Talairach^50^ N27 brain using PySurfer (https://github.com/nipy/PySurfer).

We hypothesized that geodesic social distance would be spontaneously encoded, given the importance of this information for determining self-relevance. One’s immediate social ties are obviously most self-relevant. Given the importance of reputation management for human behavior^12^, individuals two “degrees away” may be relatively important to identify and monitor: Negative interactions with such individuals could damage relationships with one’s direct connections. Similarly, sharing mutual friends may enhance trust, given the potential reputation costs of bad behavior^5^. As social distance between people increases, their relevance to each other decreases. We predicted that social distance-related information would be carried in the lateral superior temporal cortex (STC) and inferior parietal lobule (IPL), as well as the medial prefrontal cortex (MPFC), given past research implicating these regions in encoding social distance^13^ and self-relevance more generally^14^.

Social distance was reliably signaled in a large cluster centered in the lateral posterior STC and extending inferiorly throughout posterior lateral temporal cortex (LTC), and superiorly to the anterior aspect of the IPL (see Fig. 4; Table S1). Past research demonstrated that multi-voxel response patterns in this region encode egocentric spatial and abstract (e.g., social) distances when explicitly judging^14^ or mentally navigating^15^ such distances^14^; the current findings suggest that this region also encodes egocentric distances spontaneously (i.e., in the absence of any explicit distance task). Thus, when encountering a familiar individual, knowledge of agent-to-agent relationships appear to be spontaneously retrieved, such that representations of other people in this region are organized in terms of whether someone is a friend, a friend-of-a-friend, or farther removed from oneself in social ties. It has been suggested that some regions within posterior parietal cortex, such as the anterior IPL, which have well-established roles in representing and navigating physical space, analogously represent more abstract relationships (e.g., social ties between agents)^16^,^17^. The current results suggest that when encountering familiar individuals, humans may spontaneously retrieve knowledge of where they are located, relative to oneself, in a mental map of “social space”.

Although the LTC and IPL regions that carried information about social distance here have previously implicated in encoding social distance^13^,^14^, some regions previously implicated in signaling social distance were not implicated in the current study. For instance, previous research has implicated MPFC in distinguishing friends from strangers^13^, and a recent study implicated the hippocampus and posterior cingulate cortex (PCC) in tracking social distances between participants and characters in an interactive game^18^. Differences between the current results and those observed in previous investigations likely reflect differences in data analytic approaches and in how social distance has been operationalized. In the current study, participants only saw personally familiar individuals, and social distance was operationalized in terms of geodesic distance in their real-world social network. In previous neuroimaging studies, the term “social distance” has been operationalized in widely varying ways, such as the presence of social ties^13^, the strength of social ties^14^, and distance from oneself in a two-dimensional (affiliation x status) representational space^18^. Given that these variables likely have differential consequences for social cognition and behavior, it is not surprising that they are encoded by at least partially distinct neural substrates.

Whereas social distance is inherently relative to the perceiver, other aspects of familiar others’ social network positions, such as the degree to which one “bridges” different areas of the network and the number of friends someone has, are increasingly thought to be largely stable, possibly heritable, dispositional tendencies that shape social behavior^19^,^20^. Therefore, we hypothesized that EC and constraint would be encoded in brain regions involved in encoding others’ traits and behavioral tendencies more generally, such as the MPFC, which is widely implicated in inferring and encoding person knowledge^21^ and in integrating knowledge of personality traits in order to signal individual identity^22^.

Information about EC was reliably carried in brain regions that encode individual identity when imagining others’ actions^22^ (i.e., MPFC) and viewing faces^23,24^ (e.g., temporal pole; fusiform gyrus; see Fig. 4 and Table S2), suggesting that sociometric status may comprise a dimension of meaning for organizing mental representations of others. EC was also encoded in medial parietal cortex (precuneus, PCC), a region previously been shown to encode extraversion^22^, which is modestly correlated with EC^25^, suggesting that this region may encode dispositional tendencies common to both extraversion and EC. In addition, recent work has also shown that the medial parietal cortex, as well as other regions involved in inferring others’ mental states, intentions, and traits (e.g., MPFC; temporoparietal junction), responds preferentially to well-liked individuals, which is thought to reflect perceivers being preferentially motivated to understand the internal states of popular others^26^. The current findings are consistent with the notion that brain regions that represent others’ internal states and behavioral tendencies (e.g., PCC, MPFC) track sociometric status, and suggest that like other facets of social status (e.g., dominance^27^, prestige^28^), EC may modulate attention to the internal states of others. Future behavioral studies should directly test the impact of EC on social attention.

Information about EC was also reliably carried in unexpected regions, such as extrastriate visual cortex (EVC). This result is unlikely to be due to low-level visual characteristics of stimuli, as each participant had a unique stimulus set, and because videos corresponding to each individual in each stimulus set were horizontally mirrored on half of trials (see Methods). However, this finding may nonetheless reflect the effects of social status in terms of social ties on visual attention. People tend to preferentially orient toward high-status individuals and to the loci of their attention, presumably to obtain behaviorally relevant information about our surroundings^28-30^. Given that EC is reliably signaled in EVC response patterns, future research should test if visual attention is also preferentially allocated to central actors in one’s social network.

EC-based RDMs were also significantly related to neural RDMs in brain areas previously implicated in evaluating social status in terms of dominance, prestige, and morality, such as the ventral MPFC and ventrolateral prefrontal cortex (VLPFC)^31-33^. The involvement of the ventral MPFC in social status encoding has been suggested to reflect a more general role in assessing the value of stimuli^32^, whereas the VLPFC has been suggested to encode social status in order to appropriately modulate behavioral responding, which is thought to be a primary function of status cues^31^. We suggest that these regions likely encode EC for similar reasons, as high EC individuals have high behavioral relevance and value as social partners. For example, individuals connected to well-connected others may be protected from mistreatment because they are more likely to be defended by others, who themselves are more likely to be defended. Less risk is associated with wronging a low EC individual, given that low EC individuals have little influence on the spread of information and other resources^3^.

The current results suggest that when encountering a familiar individual, the degree to which that individual is well-connected to well-connected others shapes processes related to valuation, behavioral modulation, attention, and encoding others’ internal states, dispositional characteristics, and identities. Many of these findings echo the known effects of other dimensions of social status (e.g., status conferred by dominance). Although a great deal of past psychological and neuroimaging research on social status has focused on physical dominance, we note that overt physical violence is relatively rare in contemporary human groups^34^ and that social support and reputation management are central to everyday human life^12^. Social power in such groups may be relatively less contingent on individual strength and physical aggression, and more dependent on group dynamics and affiliative relationship maintenance. Thus, sociometric status is likely especially relevant to modern humans, and merits further attention in social perception and neuroscience research.

In addition to social distance and EC, diverse aspects of social cognition and behavior (e.g., deciding how to effectively seek or spread information; trust decisions) would benefit from encoding network constraint. Low constraint individuals can broker the flow of information between groups, and thus, exert a disproportionate influence on the flow of ideas and resources^4^. Additionally, individuals in relatively “closed” local networks, characterized by high constraint, suffer greater reputation costs for bad behavior; correspondingly, constraint can foster trust and cooperation^4^. Given the dearth of previous research investigating the perception of constraint, we made no specific predictions about which brain regions would be involved in encoding this facet of social network position.

Large clusters spanning both right and left lateral STC carried information about constraint (Table S3), as did a smaller cluster in the supplementary motor area (SMA). Although the lateral STC and SMA are implicated in biological motion processing^35^ and action understanding^36^, respectively, this finding was not attributable to the amount of movement in videos (see SI). A perceiver’s knowledge of the network constraint of an individual, or of associated dispositions, may impact how that perceiver attends to that individual’s movements. For example, because brokers may be perceived as exceptionally charismatic or interesting (e.g., because they often serve as sources of novel information or opportunities^4^), they may command differential amounts of attention to their expressions and gestures. Brokers may also differ in the amount of social meaning carried in their facial and bodily movements (e.g., fidgeting aimlessly vs. using movement to express oneself coherently). The latter explanation would be consistent with evidence that the STS responds to the social meaning, rather than amount, of movement in dynamic displays^37^. Future studies could arbitrate between these hypotheses by testing if strangers are able to differentiate between individuals high and low in constraint based on their observed movements. If so, this would suggest that network constraint is encoded in lateral STC because this aspect of social network position is apparent in how individuals carry themselves. If not, this would be consistent with the interpretation that perceivers’ knowledge of an individual’s network constraint, or of qualities related to this aspect of social network position, influences how perceivers attend to that individual’s expressions, gestures, and bodily movements.

After scanning, participants were asked about their perceptions of each SNA-derived metric of interest for each individual in their stimulus set (see SI). This allowed us to test the accuracy of participants’ perceptions of others’ social network positions, and to evaluate how well participants’ perceptions matched the data used to construct their stimulus sets. Post-scan ratings indicated that participants’ explicit perceptions of the social network positions of the individuals in their stimulus sets closely matched reality. Veridical constraint had a significant effect on perceived constraint (*ß*=19.44 *SE*=2.01, *p* < .0001), and veridical EC had a significant effect on perceived EC (*ß*=14.95, *SE*=0.93, *p* < .0001). Further, subjective ratings of social closeness (*ß*=-31.00, *SE*=1.62, *p* < .0001), proportion of social time spent together (*ß*=-22.74, *SE*=1.84, *p* < .0001), and frequency of discussions (*ß*=-33.77, *SE*=1.89, *p* < .0001) varied as a function of geodesic network distance (see Methods and Fig. 5).

**Figure 5.**
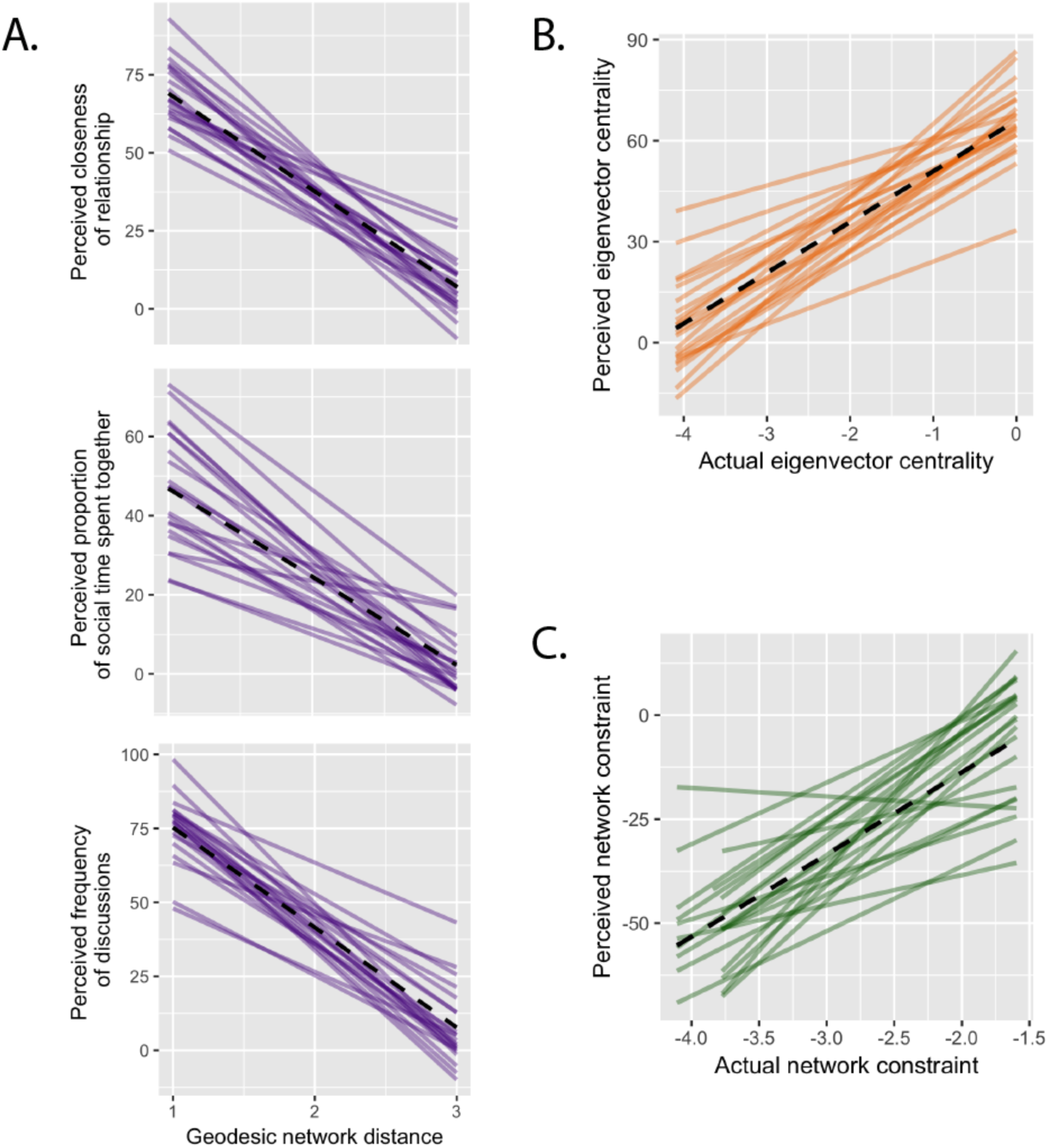
Associations between perceived and actual social network characteristics. Black dashed lines depict the relationships between perceived and actual social network characteristics across all participants (fit using an ordinary least squares linear model). Solid purple, orange and green lines depict these relationships for each subject for social distance, eigenvector centrality, and constraint, respectively. **(A)** Neuroimaging study participants’ subjective ratings of social closeness, proportion of social time spent together, and frequency of discussions with the individuals in their stimulus sets varied according to geodesic network distance from them in the network (all *p’s* < .0001, see main text). **(B)** Participants’ estimates of the eigenvector centrality of the individuals in their stimulus sets were closely related to those individuals’ actual eigenvector centralities (*p* < .0001, see main text). **(C)** Participants’ estimates of the network constraint of individuals in their stimulus sets were also associated with the actual constraint of those individuals’ positions in the social network (*p* < .0001, see main text). As described in the main text, self-report data was obtained after scanning; network constraint and eigenvector centrality were log-transformed prior to plotting and analysis to alleviate skew. Perceived network constraint ratings were multiplied by −1 prior to plotting because the relevant question asked participants to rate perceived brokerage (which is inversely related to network constraint). Analyses of behavioral ratings were conducted using linear mixed models that included bysubject random slopes and intercepts.

Although participants had consciously accessible knowledge of the social network position characteristics studied here (Fig. 5), the task used in the fMRI study (a one-back memory task) did not require participants to retrieve social relationship knowledge, and from participants’ perspectives, the social network questionnaire and fMRI study were ostensibly unrelated. Nevertheless, up to 40% of the variance in similarity structures of local fMRI responses to personally familiar others could be explained merely by characteristics of those individuals’ positions in the perceiver’s social network (Fig. 4b). These findings are consistent with behavioral evidence that humans spontaneously activate knowledge about other people upon encountering them in order to inform cognition and behavior^7^,^8^, and suggest that humans spontaneously activate complex knowledge about other people’s positions in their social networks when viewing them. These findings are consistent with psychologists’ mounting appreciation for the importance of both direct and indirect relationship knowledge to everyday cognition and behavior. Everyday interactions are influenced not only by information that would be available to any observer, but also by patterns of personal and third-party relationships. By adopting an interdisciplinary approach combining theory and methods from neuroscience, psychology, and SNA, we can begin to uncover a deeper understanding of how the human brain negotiates the intricacies of everyday social life.

## Methods

### Part 1: Social network characterization

#### Participants

Participants in Part 1 of the study were 275 first-year Masters of Business Administration (MBA) students at a private university in the United States who participated as part of their coursework on leadership (91 females; 184 males). The total class size was 277 students; two students failed to complete the questionnaire (i.e., response rate=99.3%). All procedures were completed in accordance with the standards of the Dartmouth Committee for the Protection of Human Subjects.

#### Social network characterization

To characterize the social network of all first-year students, an online social network survey was administered. Participants followed an emailed link to the study website where they responded to a survey designed to assess their position in the social network of first-year students in their academic program (see SI). The survey question was adapted from Burt^38^ and has been previously used in the modified form used here^25,39^. It read, *“Consider the people with whom you like to spend your free time. Since you arrived at [institution name], who are the classmates you have been with most often for informal social activities, such as going out to lunch, dinner, drinks, films, visiting one another’s homes, and so on?”*

A roster-based name generator was used to avoid inadequate or biased recall. Classmates’ names were listed in four columns, with one column corresponding to each section of students in the MBA program. Names were listed alphabetically within section. Participants indicated the presence of a social tie with an individual by placing a checkmark next to his or her name. Participants could indicate any number of social ties, and had no time limit for responding.

Social network analysis was performed using the R package igraph^40,41^. Three social network-derived metrics were extracted for each node: constraint, EC and geodesic distance from each classmate, as described in greater detail below.

#### Constraint

The constraint of actor *i* is given by the following equation, where *P_ij_* corresponds to the proportion of *i*’s direct social ties accounted for by his/her tie to actor *j*. The inner summation approximates the indirect constraint imposed on *i* by other actors, *q*, who are socially connected to both *i* and *j* (i.e. mutual friends of *i* and *j*):

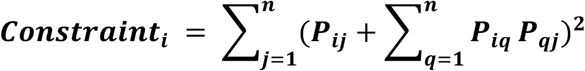

An unweighted, undirected graph was used to estimate constraint; i.e., the presence of any social tie, irrespective of its direction, was used to compute the constraint of each node. Constraint is an inverse measure of network brokerage.

#### EC

A graph consisting of nodes connected by edges can be characterized by an adjacency matrix *A*, populated by elements such that *a_ij_* = 1 if nodes *i* and *j* are directly connected, and *a_ij_* = 0 if these nodes are not connected. The EC of each node is given by the eigenvector of *A* in which all elements are positive. The requirement that all elements of the eigenvector must be positive yields a unique eigenvector solution (i.e., that corresponding to the greatest eigenvalue). Here, when computing EC, the directionality of the graph was preserved; in the event of asymmetric relationships, only incoming, rather than outgoing, ties were used to compute EC.

#### Social distance

Geodesic social distance refers to the smallest number of intermediary social ties required to connect two individuals in a network. Individuals who a participant named as friends have a distance of one from him/her. Individuals whom a participant’s friends named as friends (but who were not named as friends by the participant) have a distance of two from the participant. Individuals who were named as friends by classmates at a distance of two from the participant (but not by the participant or his/her friends) have a distance of three, and so on.

### Part 2: Neuroimaging study

#### Participants

A subset of individuals who had completed Part 1 participated in a subsequent neuroimaging experiment. Participants were informed during class about the opportunity to participate in an fMRI study that was ostensibly unrelated to the online questionnaire in Part 1, and that they would receive $20/hour as compensation and images of their brains. All participants were right-handed, fluent in English, and had normal or corrected-to-normal vision. Participants provided informed consent in accordance with the policies of the Dartmouth College Committee for the Protection of Human Subjects. Twenty-four participants (12 females) completed the fMRI study. The sample size was chosen based on previous fMRI studies using similar paradigms and RSA methods^11,42^. One participant was excluded due to image artifact, and two were excluded because they scored less than 65% correct on the one-back memory task used in the scanner (this threshold was based on what has been used previously in similar studies^43^). Consequently, we analyzed data from 21 participants (10 females, aged 25-33, *M*=27.95, *SD*=2.16). As a within-subjects design involving no group allocation was used, blinding investigators to between-subjects conditions and random assignment of participants to conditions were not applicable.

#### Image acquisition

Participants were scanned at the Dartmouth Brain Imaging Center using a 3T Philips Achieva Intera scanner with a 32-channel head coil. An echo-planar sequence (35 ms TE; 2000 ms TR; 3.0 mm x 3 .0 x 3.0 mm resolution; 80 x 80 matrix size; 240 x 240 mm FOV; 35 interleaved transverse slices with no gap; 3.0 mm slice thickness) was used to acquire functional images. Functional runs consisted of 180 dynamic scans, for a total acquisition time of 360 s per run. A high-resolution T1-weighted anatomical scan was acquired for each participant (8.2 s TR; 3.7 ms TE; 240 x 187 FOV; 0.938 mm x 0.938 mm x 1.0 mm resolution) at the end of the scanning session. Foam padding was placed around subjects’ heads to minimize motion.

#### Stimuli

Each participant’s customized stimulus set consisted of short videos of four individuals at each of three geodesic distances (i.e., one, two, and three) from the participant in the social network of first-year MBA students. The two highest and lowest EC individuals at each social distance were included in the stimulus set (Fig. 2).

The videos used as stimuli consisted of individuals introducing themselves (e.g., “Hi my name is [first name], and you can call me [first name/nick name]”). A video of this kind was made involving each student at the beginning of the academic year as a resource for other students and faculty. Videos were truncated to 2 s, beginning when the subject began to say the word, “Hi,” and were presented without sound. Prior to entering the fMRI scanner, participants were shown each video with sound to familiarize themselves with the stimuli.

#### fMRI paradigm

The fMRI study consisted of 10 runs and followed a rapid event-related design with an inter-trial interval consisting of 4 s of fixation (Fig. 2c). Four null events, each consisting of an additional 2 s of fixation, were randomly inserted into each run. In each run, four repetitions of 14 event categories (12 identities; 1 null event; 1 catch trial) were pseudo-randomized such that there were no consecutive repeats of the same category. Horizontal mirroring was randomly applied to half the presentations of each stimulus within each run to reduce similarities within identities due to local low-level visual features. Catch trials involved seeing the same stimulus at the same mirroring level as the immediately previous stimulus (or two trials back if a catch trial followed a null event). Participants were instructed to press a button when an identical video was presented twice in a row (i.e., for catch trials).

#### Post-scan questionnaire

After scanning, participants were asked about their subjective perceptions of each social network metric of interest for each individual in their stimulus set, as well as questions assessing tie strength (see SI). Because the constraint question asked about brokerage (i.e., which individuals were low in network constraint), responses to this item were multiplied by −1. To alleviate skew in the network data, eigenvector centralities and network constraint values were log-transformed prior to analysis.

The correspondence between participants’ post-scan ratings and the social network position characteristics of the individuals in their stimulus sets was assessed using linear mixed models using the R^40^ package lme4^44^. For each of the five questions (see SI), a model was constructed with participants’ ratings as the dependent measure and the relevant social network position characteristic as a fixed effect, as well as random intercepts and slopes for each participant. To test the significance of the relationship between participants’ ratings and social network data, p-values were computed using Satterthwaite’s approximation for degrees of freedom^45^ as implemented in lmerTest^46^.

#### fMRI data preprocessing

For fMRI data analysis, data were preprocessed and average voxel-wise hemodynamic responses to each identity were estimated using AFNI^47^. Pre-processing steps included applying AFNI’s 3dDespike function to remove transient, extreme values in the signal not attributable to biological phenomena, slice-timing correction to correct for interleaved slice acquisition order, alignment of the last volume of the final run to the high-resolution anatomical scan, registration of all functional volumes to the anatomical-aligned final functional volume using a 6-parameter 3-D motion correction algorithm, spatial smoothing using a 4-mm full-width at half-maximum Gaussian kernel, and scaling each voxel time series to have a mean amplitude of 100. Prior to regression, consecutive volumes where the Euclidean norm of the derivatives of the motion parameters exceeded 0.3 mm were excluded from further analysis, as were volumes in which more than 10% of brain voxels were identified as outliers by the AFNI program 3dToutcount.

Parameter estimates were extracted for each voxel using a GLM that consisted of gamma-variate convolved regressors for each of 13 predictors (one for each of the 12 identities in the participant’s stimulus set; one for catch trials), as well as 12 regressors for each of the six demeaned motion parameters extracted during volume registration and their derivatives, and three regressors for linear, quadratic, and cubic signal drifts within each run. This procedure removed variance caused by regressors of no interest, and resulted in an estimate of the response of each voxel to each trial type.

#### GLM decomposition searchlight

Using PyMVPA^48^ and SciPy^49^, a GLM decomposition searchlight^11^ was performed within each participant’s data. A sphere (radius=3 voxels) was moved throughout each participant’s brain. At each point in the brain, the local distributed patterns of neural responses to each person in the stimulus set were extracted within a sphere centered on that point, and the pairwise correlation distances between them were calculated to construct a local neural RDM (Fig. 3a-c), which was decomposed into a weighted combination of predictor RDMs using ordinary least squares (OLS) regression (Fig. 3d). There were three predictor RDMs, one corresponding to each social network position metric of interest. Predictor RDMs were constructed by taking the Euclidean distance (i.e., the absolute value of the numerical difference) between the relevant social network position metrics for each possible pair of identities within each participant’s stimulus set. Each predictor RDM for each participant was then z-scored. Next, for each RDM (e.g., the EC-based RDM for a given participant), the variance accounted for by the remaining two predictor RDMs (e.g., the social distance and constraint-based RDMs for that participant) was removed using OLS regression. Thus, the resultant predictor RDMs were made orthogonal to one another prior to performing the GLM decomposition searchlight.

At each searchlight center (i.e., at each voxel), the GLM decomposition procedure yielded a *β* value corresponding to each social network derived metric of interest, as well as an *R*^2^ value corresponding to how much the variance in the similarity structure of local neural response patterns could be explained by the social network positions of the individuals comprising a given participant’s stimulus set.

#### Group analysis

Each subject’s maps of regression coefficients and *R^2^* values were transformed to standard (Talairach^50^) space using AFNI: Anatomical scans were linearly aligned to the Talairach^50^ template using the @auto_tlrc algorithm in AFNI, and the same transform was used to align each participant’s searchlight results to standard space prior to group analysis. To identify areas that reliably contained information about each specific aspect of social network position across participants, the regression coefficients for each social network position-derived RDM were tested against 0 across participants using one-tailed one-sample *t*-tests. More specifically, FSL’s randomise^51^,^52^ program was used to perform permutation tests and to generate a null distribution of cluster masses for multiple comparisons correction (cluster-forming threshold: *p* < .01, two-tailed; 5,000 permutations; 10-mm variance smoothing). All reported results have been thresholded at a family-wise error rate of 5%.

#### Data and code availability

The data that support the findings of this study are available from the corresponding author upon request. The code used for the analyses also is available upon request.

## Acknowledgements

This work was supported by a graduate fellowship from the Neukom Institute for Computational Science and a Dartmouth Graduate Alumni Research Award. The authors wish to thank Will Haslett for assistance with the optical flow analysis.

## Author Contributions

CP, AMK and TW conceived of and designed the study. CP and AMK collected the data. CP analyzed the data. CP, AMK, and TW wrote the paper.

## Supplementary Information

### I. Optical flow analysis

To quantify the amount of movement within each video clip used as a stimulus in the neuroimaging experiment, the average optical flow (i.e., the pattern of apparent motion between consecutive video frames) was computed for each video that was shown in the fMRI study. Given that the videos used as stimuli were recorded by a stable camera against a plain, static background, optical flow estimates for these videos capture of the amount that each individual moved his or her facial features and head in the video clip. Farneback’s algorithm for motion estimation^1^ as implemented in OpenCV^2^ was used to estimate the average magnitude of optical flow in each video. This method extracts a pixel-wise motion vector for each pair of sequential frames in which each pixel is characterized by a magnitude and a direction. To estimate the magnitude of motion within each frame pair, the magnitude values (without respect to direction) were summed across pixels. To compute the mean magnitude of optical flow for a given video, the motion magnitude estimates were averaged across frames within that video.

In order to test whether or not individual differences in network constraint are related to movement in the videos used as stimuli, the correlation between network constraint and average motion magnitude was assessed among the 88 individuals whose videos were used as stimuli in the fMRI study. Given that distributions of both variables were highly skewed, data were log-transformed prior to analysis. The results of this procedure suggest that in the stimuli used in the current study, network constraint and amount of movement were not significantly correlated, *r* = −0.12, *p* = 0.28 (see Fig. S1).

### II. Post-scan questionnaire

Participants performed the post-scan questionnaire on a 13” MacBook laptop using Psychopy^3^. Participants first viewed an instruction screen that read, *“Now you will see the same people who you saw in the scanner. You will be asked questions about each person. These questions relate only to this person’s interactions within the [institution name] MBA cohort. We understand that people have many social circles that they participate in (perhaps including family, friends outside of [the institution], other contacts, etc.). For these questions, please just consider interactions within the MBA cohort. You will be presented with a continuous rating scale for each question. You can choose any point along the continuum to respond. Press any key to continue.”* During the survey, videos of the 12 individuals from the participant’s stimulus set were presented in a random order. Participants responded to all questions about a given individual sequentially, and the same video that had played in the scanner repeated on a loop (without sound) above the question text and response scale (see Fig. S2).

Participants were presented with questions concerning lay definitions of eigenvector centrality (*“In social network analysis, scientists assess a construct that measures how many friends a person has, and how many friends a person’s friends have. How would you rate this person on this construct?”* Responses ranged from *“Low (few friends who have few friends)”* to *“High (many friends who have many friends)”*) and constraint (*“Social network analysts also assess a construct called ‘brokerage’ that measures how much a person connects groups of people who wouldn’t otherwise be connected. Using this definition, how high is this individual in ‘brokerage’?”* Responses ranged from *“Low (this person never connects distinct groups of people”* to *“High (This person often connects distinct groups of people)”*). Responses to the item assessing brokerage were reverse scored in order to estimate perceived network constraint.

Participants were also presented with the name generator that had originally been used to construct the network (*“Consider the people with whom you like to spend your free time. During the last month, is this one of the classmates who you have been with most often for informal friendship activities, such as going out to lunch, dinner, drinks, films, visiting one another’s homes, and so on?”* Responses ranged on a continuum from *“None of my social activities in the past month have included this person”* to *“All of my social activities in the past month have included this person”*), as well as questions designed to assess tie strength (*“How close are you with this person?”* Responses ranged from *“Distant”* to *“Less than close”* to *“Close”* to *“Especially Close”*) and frequency of interactions (*“On average, how often do you talk to this person (any social or business discussion*)*?”* Responses ranged from *“Less often”* to *“Monthly”* to *“Weekly”* to *“Daily”*).

---

**Figure S1.**
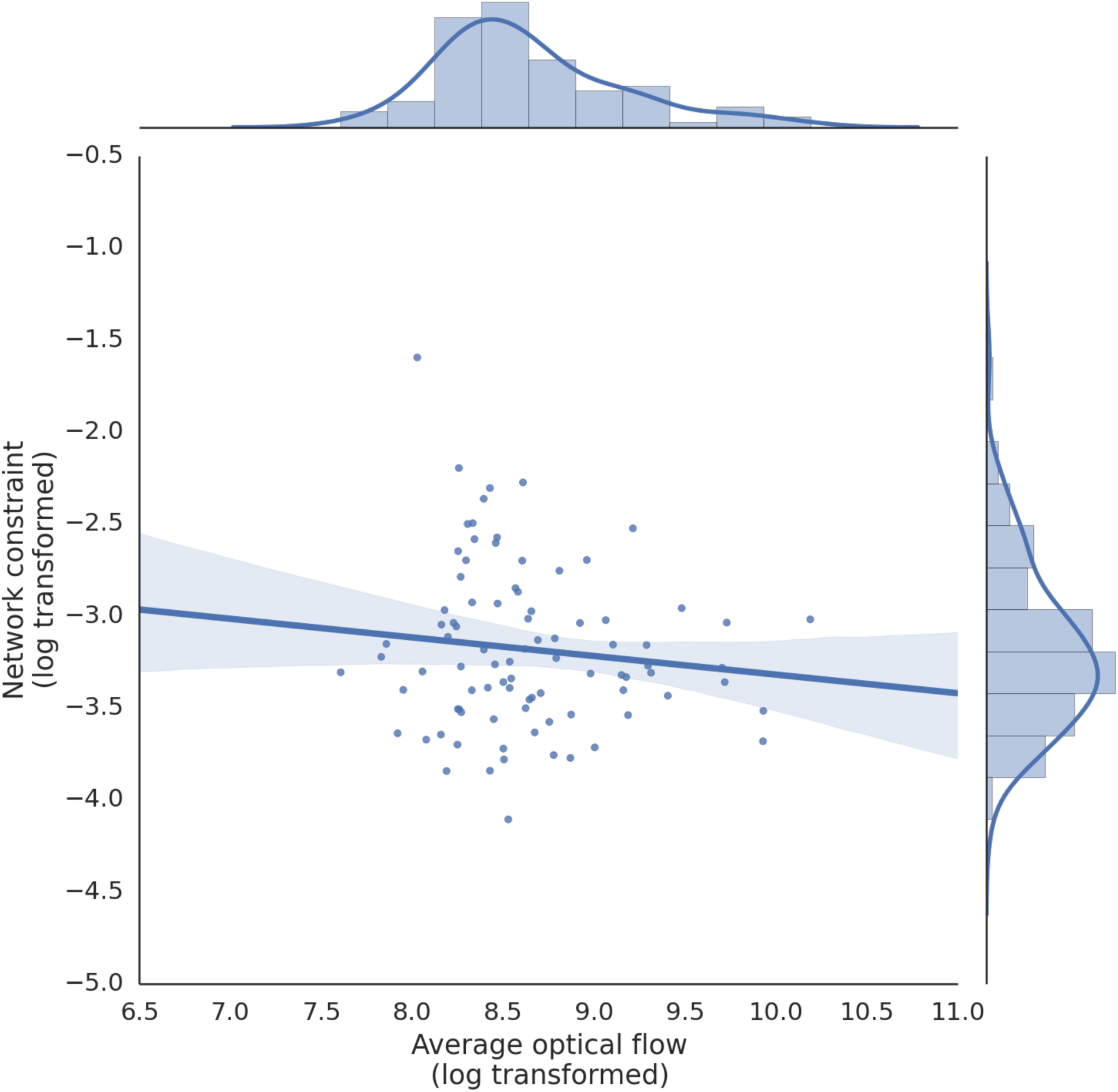
Relationship between network constraint and movement during videos. The amount of movement of the 88 individuals whose videos were used as stimuli was not significantly related to the constraint characterizing those individuals’ positions in the social network of first-year MBA students, *r* = −0.12, *p* = 0.28.

**Figure S2.**
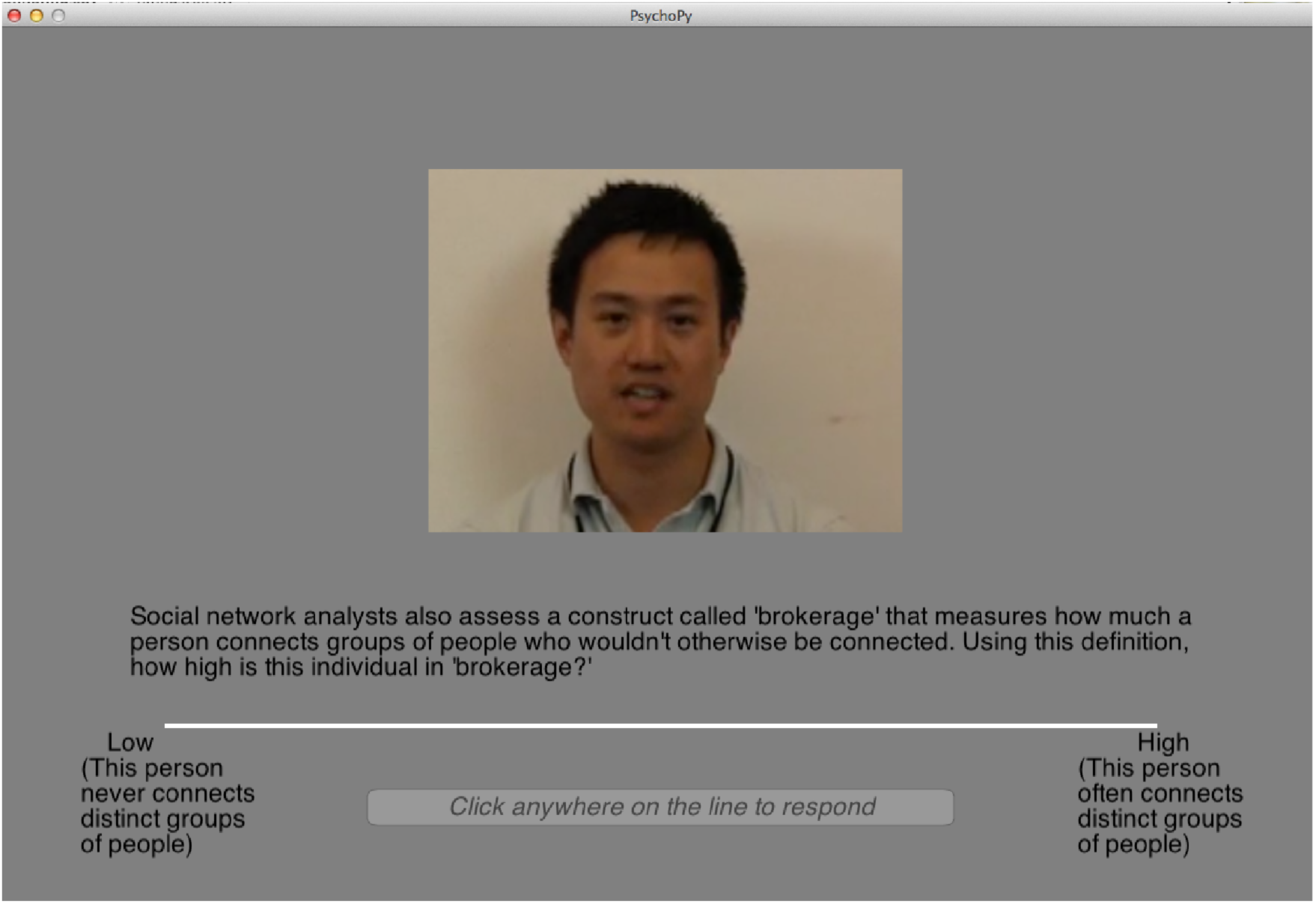
Post-scan questionnaire. Following scanning, participants responded to questions about their subjective perception of each aspect of social network position of interest for each individual in their stimulus set. A screenshot of the question corresponding to network constraint (reverse-scored) is shown.

**Table S1:** Brain regions where local neural information content is associated with the social distance of the individuals being viewed.

| Hemi | Size<br>(mm<sup>3</sup>) | COG<br>x | COG<br>y | COG<br>z | Location |
| --- | --- | --- | --- | --- | --- |
| R | 4,397 | 49.6 | -46.6 | 7.6 | IPL (SMG), STG, STS, MTG |

Hemi = hemisphere; COG = center of gravity; L = left; R = right; IPL = inferior parietal lobule; SMG = supramarginal gyrus; STG= superior temporal gyrus; STS = superior temporal sulcus; MTG = middle temporal gyrus. All reported results are significant at a statistical threshold of $p < .05$ , FWE-corrected. All coordinates are in Talairach space.

**Table S2:** Brain regions where local neural information content is associated with the eigenvector centrality of the individuals being viewed.

| Hemi | Size<br>(mm<sup>3</sup>) | COG<br>x | COG<br>y | COG<br>z | Location |
| --- | --- | --- | --- | --- | --- |
| L | 24,483 | -42.4 | -18.5 | 33.6 | IPL, IFG, Ins., pre-central gyrus |
| R | 8,768 | 21.7 | 26.8 | -4.9 | MPFC, IFG, aIns., ant. PHG, TP |
| L | 7,716 | -32.7 | 12.9 | -6.1 | aIns., IFG |
| L, R | 7,552 | -9.4 | -41.8 | 40.4 | PCC, precuneus |
| R | 6,802 | 20.4 | -48.0 | -2.8 | PHG, LG, FG |
| R | 6,065 | 48.0 | -29.4 | 38.4 | IPL, precuneus, post-central gyrus |
| L, R | 5,233 | -0.8 | -44.4 | 65.4 | Precuneus, post-central gyrus |
| L, R | 4,961 | -0.6 | -83.3 | 28.5 | EVC |

Hemi = hemisphere; COG = center of gravity; L = left; R = right; a = anterior; IPL = inferior parietal lobule; IFG = inferior frontal gyrus; Ins. = insula; MPFC = medial prefrontal cortex; PHG = parahippocampal gyrus; TP = temporal pole; PCC = posterior cingulate cortex; LG = lingual gyrus; FG = fusiform gyrus; EVC = extrastriate visual cortex. All reported results are significant at a statistical threshold of $p < .05$ , FWE-corrected. All coordinates are in Talairach space.

**Table S3:**
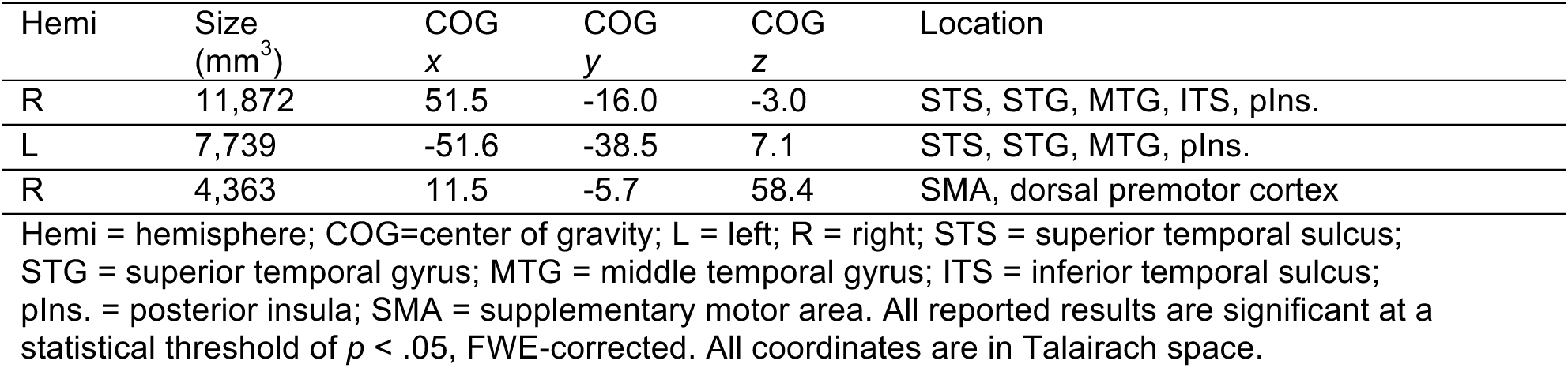
Brain regions where local neural information content is associated with the constraint of the individuals being viewed.

